# Generation of cloned sheep lacking galactose-α1,3-galactose and N-glycolylneuraminic acid antigens

**DOI:** 10.1101/2025.10.31.684537

**Authors:** Sarah J Appleby, Lisanne M Fermin, Zachariah L McLean, Stephanie Delaney, Stefan Wagner, Cecilia Di Genova, Jingwei Wei, Fanli Meng, David N Wells, Björn Oback

## Abstract

Livestock have long been regarded as a potential source of donor organs to alleviate the global organ shortage for transplantation. Sheep have a similar physiology and anatomy to humans, providing the standard model for demonstrating biocompatibility and performance of biological heart valves to obtain regulatory approval for their use in transplantation. Like most mammals, sheep cells contain two well-characterized carbohydrate epitopes, galactose-α1,3-galactose (α-Gal) and *N*-glycolylneuraminic acid (Neu5Gc), which are absent in humans. Formation of these xenoantigens is catalysed by two enzymes, namely α(1,3) galactosyltransferase (GGTA1) and CMP-Neu5Gc hydroxylase (CMAH), respectively. Towards generating new sheep models, we used Cas9-mediated genome editing in both male and female ovine fetal fibroblasts to knockout (KO) both alleles of *CMAH* and *GGTA1*. Selected double KO (DKO) fibroblast strains were used for somatic cell transfer cloning to produce blastocysts that were transferred into gestational surrogate ewes. Following transfer of 128 male and 40 female cloned blastocysts, 6 male and 8 female lambs were born, however, none of the males survived. Molecular analyses of cells from the five surviving ewes confirmed their compound heterozygous state, resulting in a functional DKO phenotype and subsequent immune rejection of embryonic tissues during natural breeding with wild-type rams. This new DKO sheep model better mimics the human immune status, offering greater physiological relevance for preclinical testing of biological heart valves, alleviation of red meat allergy syndrome due to the presence of dietary xenoantigens, and an alternative source of donor organs that may be culturally more widely accepted than pigs.

## Introduction

The global gap between short supply and high demand of donor organs persists, limiting the lifesaving potential of organ transplantation. With an ever-increasing demand for replacement organs, livestock have long been regarded as potential donors for transplantation into humans. However, xenotransplantation is hampered by the immediate host immune response towards donor organs due to the presence of naturally formed cytotoxic antibodies.

The major xenoantigen galactose-α1,3-galactose (α-Gal) is formed by the addition of galactose to *N*-acetyllactosamine by the enzyme α1,3-galactosyl transferase 1 (GGTA1). α-Gal is present in all mammals, except Old World monkeys, apes, and humans. Due to the abundance of α-Gal in their environment, especially gut bacteria and red meat, Old World primates develop large amounts of antibodies directed towards α-Gal. These antibodies are the most abundant natural antibody in humans [1] and are responsible for 70–90% of the human antibodies that bind to pig cells in xenotransplantation experiments [2].

Despite α-Gal being responsible for the majority of naturally formed antibodies causing hyperacute rejection of xenografts, non-Gal epitopes are of concern once α-Gal is removed. The sialic acid *N*-glycolylneuraminic acid (Neu5Gc) was identified as the next crucial xenoantigen, accounting for the majority of non-Gal antibodies [3]. Neu5Gc formation is catalysed by the enzyme cytidine monophosphate-Neu5Ac hydroxylase (CMAH), and is absent in humans, New World monkeys, and a small number of other species through convergent evolution [4]. Continued pursuit of xenoantigens has further led to the identification of DBA-reactive glycans (also named Sd(a) antigen) produced by β-1,4-*N*-acetylgalactosaminyl transferase 2 (β4GalNT2), another glycan associated with xenograft injury induced by highly specific circulating human antibodies [5].

Gene editing technologies generated single *GGTA1* knockout (KO) pigs [6] to evaluate their use for xenotransplantation. Following the initial generation of α-Gal absent *GGTA1*^-/-^ pigs, various tissues from α-Gal negative pigs have been transplanted into non-human primates to evaluate survival.

Following transplantation into baboons, hearts and kidneys from α-Gal negative pigs survived 59–179 days [7] and 8–16 days [8], respectively.

Removal of both α-Gal and Neu5Gc in double KO (DKO) *GGTA1^-/-^ CMAH^-/-^* pigs was demonstrated to lower the immune response from human sera compared to *GGTA1* KO alone [9]. Further study showed that the human immune response elicited by *GGTA1*/*CMAH* KO pig serum was also lower than that elicited by serum from chimpanzee, the species with greatest xenotransplant survival in early studies [10].

Most large animal studies for use in human therapeutics are focused on pigs, due to similarities in physiology and diet [11]. Kidney size (relative to bodyweight) and physiology, such as nephron number, is similar between pigs, sheep, and humans. However, sheep are more similar to humans regarding total kidney weight, blood flow, and timing of kidney development. Since sheep kidneys are slightly smaller than humans, but have similar blood flow, their kidney perfusion rate is higher than in humans and pigs [12,13].

First attempts to clone *GGTA1* KO sheep did not produce live animals, possibly due to the prolonged *in vitro* culture required for isolating cells targeted by homologous recombination [14]. Subsequently, viable *GGTA1* null sheep were produced via Cas9-mediated editing and somatic cell transfer (SCT) cloning, reviving interest in this species as a preclinical model, particularly for cardiovascular applications such as heart valve replacement [15].

Sheep serve as the primary experimental and regulatory testing model for the development and approval of new mechanical and biological heart valves [16]. Valve implantation into sheep is used to test physiological compatibility and hemodynamic function, as well as biological responses implicated in structural valve degeneration. Despite its widespread use, the standard valve implantation sheep model is compounded by its clinical immune responses to xenoantigens. Immune rejection, mediated by anti-Gal and non-Gal circulating antibodies, may contribute substantially to age-dependent degeneration of transplanted biological heart valves. This hypothesis has not been fully testable in normal sheep because they do not produce anti-xenoantigen antibodies. Recently, GalKO sheep have been shown to produce cytotoxic anti-Gal IgG at clinically relevant levels [15]. This provides an animal model to directly test the role of the anti-gal antibody on bioprosthetic heart valve transplants, mimicking the human immune responses to α-Gal. To further refine this model, it is critical to also account for the effect of non-Gal related immune responses as an immune model for regenerative heart valve development.

Here, we used Cas9-mediated genome editing to disrupt *CMAH* and *GGTA1* simultaneously in ovine fetal fibroblasts. Edited donor cell strains were used to clone animals lacking the xenoantigens Neu5Gc and α-Gal for phenotyping. These advances align with the trend towards multi-pathway immune modulation in xenotransplantation and support the growing role of sheep as an immunologically matched and ethically acceptable species for transplantation and clinical testing.

## Methods

Experiments complied with the New Zealand Animal Welfare Act 1999 and were approved by the Ruakura Animal Ethics Committee (applications 14258 and 14693).

### Cell source and culture

Ovine fetal fibroblasts (OFFs) were previously isolated from abattoir fetuses following standard operating procedures in cattle [17]. Briefly, fetuses were washed with ethanol, skin dissected, washed briefly again in 70% ethanol, minced, and cultured in hanging drops. Male OFF3 cells were isolated from a Perendale predominant composite fetus approximately 84–98 days post conception (dpc), and female OFF4 cells from a Coopworth predominant composite fetus under 42 dpc [18]. OFFs were cultured in Dulbecco’s Modified Eagle Medium: Nutrient Mixture F-12 with GlutaMAX™ supplement (DMEM/F12; Gibco, Waltham, MA, USA) with 10% fetal bovine serum (FBS; Moregate Biotech, New Zealand) at 38°C in a humidified 5% CO_2_ incubator. Cells were passaged using 0.25% trypsin-EDTA (Gibco) for 2–5 min and centrifuging at 200 x g for 3–5 min.

### DNA extraction

Genomic DNA was isolated from OFF3 and OFF4 cells using lysis buffer (100 mM Tris pH 8, 1 mM EDTA, 0.5% (v/v) Tween-20, 0.5% (v/v) Triton X-100) containing 1 mg/mL Proteinase K (QIAGEN, Germany) and incubated at 55°C for 15 min. Proteinase K was heat inactivated by incubating for 5 min at 95°C. DNA was either used as a crude lysate or ethanol precipitated if a purified sample was desired.

### Endpoint polymerase chain reaction (PCR)

The genomic region of interest was amplified with endpoint PCR using KAPA2G (Kapa Biosystems, Wilmington, MA, USA) following the kit instructions. Briefly, the 25 µL reaction contained 0.5 µM primers and 1 µL template DNA. Cycling conditions were: initial denaturation at 95°C for 3 min; 35 cycles of 95°C for 15 s, 60°C for 15 s, and 72°C for 1 s; and final extension at 72°C for 5 min. PCR was confirmed on a 1% agarose gel with 1 Kb Plus DNA ladder (Invitrogen, Waltham, MA, USA). Primers (Table S1b) were designed using PrimerBLAST [19] for *Ovis aries* with product size 400–1000 bp. For off-target analysis, primers were designed with the off-target site at least 50 bp away from one primer that could be used for sequencing.

### Sequencing

DNA was purified from the PCR reaction using NucleoSpin Gel and PCR Clean-up kit (MACHEREYNAGEL, Germany), according to the manufacturer’s instructions, and concentration quantified using a Nanodrop (version 1000; ThermoFisher, Waltham, MA, USA). Sanger sequencing was performed by Massey University Genome Service (Massey University, New Zealand). Sequences were mapped to the reference genome in Geneious Prime (Biomatters, New Zealand) and consensus sequence generated.

### gRNA-Cas9-mediated disruption of *CMAH* and *GGTA1*

#### *CMAH* guide design

Potential target locations in sheep were chosen based on similarities to previous targeting sites in pig and the potential to disrupt the majority of the functional domains. Similarity to the reference genome was confirmed for the target regions of *CMAH* in the parental OFF3 and OFF4 cell lines using Sanger sequencing.

Once the sequence was confirmed, constructs were designed for targeting exon 5 of *CMAH*. Three gRNA were designed (gRNA1, gRNA2, and gRNA3. Sequences in Table S1a) on CRISPOR within a 202 bp section including all of exon 5. Targeting constructs were chosen that had high on-target efficiency, low off-target sites, and were in close proximity. Oligos for top and bottom strands of the gRNA sequence were ordered with appropriate overhangs for ligation into Cas9 plasmids [20]. Guide sequences had a single ‘G’ added if the sequence did not start with that nucleotide to improve transcription from the U6 promoter.

#### *CMAH* plasmid generation

*CMAH* gRNA were inserted into Cas9 plasmid PX459 (containing puromycin resistance) following the protocol from the Zhang lab [20]. Briefly, PX459 was digested with *Bbs*I-HF in Cutsmart buffer (New England Biolabs, Ipswich, MA, USA) overnight at 37°C, purified by running on an agarose gel, and DNA isolated using the NucleoSpin kit. *CMAH* gRNA oligos were annealed together (without phosphorylation) using an incubation at 95°C for 10 min followed by cooling to room temperature at 5°C min^-1^. Annealed oligos were diluted 1:200 in MilliQ H_2_O and ligated into the digested plasmid. In a 20 µL reaction, 1 µL diluted oligos were ligated into 100 ng digested plasmid using 1 µL T4 ligase in 10 µL 2X T4 ligase buffer (ligase and buffer from pGEM®-T Easy Vector kit; Promega, Madison, WI, USA) made up to full volume with MilliQ H_2_O. The reaction was incubated for either 2 h at room temperature or overnight at 4°C, before transformation into DH5α (Zymo Research, Irvine, CA, USA). Transformed bacteria were plated on LB agar with 100 µg/mL ampicillin (Sigma-Aldrich, St Louis, MO, USA) and incubated overnight at 37°C. Selected colonies were amplified in 3 or 300 mL LB (Invitrogen) with ampicillin. Plasmids were extracted from amplified bacteria using PureLink® HiPure Plasmid Filter Miniprep and Maxiprep Kits (Invitrogen), respectively. Insertion of gRNA was confirmed by performing an endpoint PCR with the top gRNA oligo as the forward primer and a primer designed to bind within the CBh promoter and run on an agarose gel. Plasmid sequence was confirmed by Sanger sequencing using a primer upstream of the gRNA insertion site.

#### *GGTA1* guide design and plasmid generation

*GGTA1* gRNA had been previously designed for targeting in goat and inserted into the Cas9 plasmid PX330. Briefly, PX330 was digested with *BbsI* in 10X NEBuffer 2.1 (New England Biolabs) for one hour at 37°C, purified by running on an agarose gel, and DNA isolated using the NucleoSpin kit. *GGTA1* gRNA oligos were phosphorylated using T4 polynucleotide kinase (Invitrogen) and annealed at 37°C for 30 minutes, followed by 95°C for 5 min and cooling to room temperature at 5°C min^-1^. Annealed oligos were diluted 1:250 in MilliQ H_2_O and ligated into 50 ng of digested plasmid using 5 units T4 ligase (Roche). The reaction was incubated for 2 h at room temperature, before transformation into DH5α (Zymo Research, Irvine, CA, USA). Transformed bacteria were plated on LB agar with 100 µg/mL ampicillin (Sigma-Aldrich, St Louis, MO, USA) and incubated overnight at 37°C. Selected colonies were amplified in LB with ampicillin. Plasmids were extracted using PureLink® HiPure Plasmid Filter Miniprep and Maxiprep Kits, respectively (Invitrogen). Insertion of gRNA was confirmed by double digestion reaction employing *BbsI* and *AgEI* restriction enzymes and run on an agarose gel. Plasmid sequence was confirmed by Sanger sequencing using a primer upstream of the gRNA insertion site. Sequence conservation between goat and sheep was confirmed using reference genomes. Sanger sequencing was performed on OFF3 and OFF4 to ensure the sequence of the most efficient guide used in goat (gRNA4; Table S1) matched the sheep cell lines.

### Transfection and editing efficiency

*CMAH* and *GGTA1* plasmids were transfected into 1.5 x 10^5^ cells using the Neon® Transfection System (ThermoFisher) with the 10 µL electrode pipette tip according to the manufacturer’s instructions. Briefly, cells were passaged, washed in 10 mL phosphate buffered saline (PBS), and resuspended at 1.5 x 10^7^ cells/mL in room temperature Re-suspension Buffer. Plasmid combinations were added to 1.5 mL Eppendorf tubes on ice; the plasmid concentration and volume did not exceed 1 µg DNA per 1 x 10^5^ cells and 10% of the total reaction volume. A total of 10 µL of resuspended cells (1.5 x 10^5^ cells) was added to the plasmid tubes and mixed briefly. Using the 10 µL Neon™ electrode pipette tip, cells were transferred to the Neon™ system and electroporated at 1,500 V with one 20 ms pulse. Cells were ejected into a prepared 4-well or 35 mm tissue culture plate with standard culture medium. Transfected cells were transiently selected with puromycin (2 µg/mL) for 48 h for PX459 uptake. Cells were grown for 7 d and genomic DNA isolated. All three *CMAH* gRNAs were screened in OFF3 and the most efficient guide was used to target OFF4.

Editing efficiency of both genes in OFF3 and OFF4 was measured using the Tracking of Indels by DEcomposition (TIDE) webtool [21]. Genomic DNA was extracted from transfected cell samples and prepared for Sanger sequencing. Sequencing files (abi) for edited sample and unedited control were uploaded to the TIDE website, with all settings left at default and an indel size of 35, unless otherwise stated. Results were accepted if R^2^ value ≥0.89.

### Strain isolation and screening

Clonal cell strains were isolated from chosen edited populations using manual selection of mitotic doublets into a 96-well plate, as described previously [22]. Cells were expanded and passaged through 48- and 24-well plates, before cryopreservation, DNA isolation, and endpoint PCR. If two bands were visualised on the agarose gel, they were cut out using a Safe Imager™ Blue Light Transilluminator (ThermoFisher) and DNA isolated separately. DNA was purified using the NucleoSpin kit. Strains were sequenced and screened on Geneious and TIDE to identify the specific mutations in both genes. Simple editing events within a sample (single indel) were identified on Geneious and confirmed with TIDE. Multiple edits (compound heterozygotes) within a sample caused two peaks at each nucleotide position in the sequence after the cut site. Indels smaller than 35 bp were identified with TIDE. Larger indels were estimated in Geneious by manually identifying the two nucleotides sequenced at each location and matching to the wild-type (WT) sequence upstream of the cut site. Strains were considered clonal if they contained a maximum of two edits at a single locus (one edit per allele). Multiple strains with identical indels were grouped into ‘mutation profiles’, indicating these strains likely came from the same original edited cell. Desirable mutation profiles contained small indels (insertions <5 bp or deletions <30 bp) that caused a frameshift (single edit per allele and indel not a multiple of 3).

Strains were assessed for integration of PX459 or PX330 into the sheep genome. Endpoint PCR was performed with primers designed to bind within the Cas9 coding sequence of the plasmid and Cas9 presence within a strain was confirmed by a band on an agarose gel. Diluted PX459 and WT DNA were used as positive and negative controls, respectively. A representative of each mutation profile for OFF3 and all OFF4 strains was assessed. The Cas9 insertion assay was run alongside a PCR for a sheep gene (*CMAH* or *GGTA1*) to confirm DNA was present from each sample. Strains positive for Cas9 were excluded from further analysis.

A biased Cas9 off-target screen was performed on strains with a desirable mutation profile. CRISPOR was used to identify the most likely off-target sites for *CMAH* gRNA1 and *GGTA1* gRNA4 used in both OFF3 and OFF4 cell lines. The top three target sites for each guide (based on CRISPOR ranking) were chosen for analysis. Genomic regions were amplified, sequenced, and compared with the WT sequence for any evidence of editing.

### SCT and embryo transfer (ET)

Sheep somatic cell cloning was carried out using a protocol published previously [18]. Briefly, donor OFF cells were synchronised via serum starvation in 0.5% FBS for 4–6 d and harvested using trypsinisation. Zona-free *in vitro* or *in vivo* matured MII oocytes were enucleated and donor cells attached in the presence of phytohemagglutinin P (L9017, Sigma-Aldrich). Couplets were fused in 270 mOsm fusion buffer without calcium with an ∼80 V/cm alternating current for automatic alignment followed by two direct current pulses of 2 kV/cm for 10 µs. Reconstructs were activated 3–4 h post fusion with ionomycin (I0634, Sigma-Aldrich)/6-dimethylaminopurine (D2629, Sigma-Aldrich). After 4 h, reconstructs were washed and cultured in groups of 10–12 embryos per 20 µL early synthetic oviduct fluid (ESOF) drop within individual micro-wells. Media was refreshed on day 2 and changed to late SOF (LSOF) on day 4. Embryos were cultured in individual 5 µL LSOF drops from day 5 onwards. Cloning runs completed in 2019 used morula aggregation in attempts to improve embryo quality and *in vivo* survival to term [18]. On day 5, morulae were broken into 2–3 small fragments and all fragments of 2 morulae were aggregated together. Blastocysts were morphologically graded on day 6 or 7 and transferred into synchronised recipient ewes or vitrified for future transfer [23].

For ET, the oestrus cycle of recipient ewes was synchronised using progesterone CIDR (EAZI-BREED™ CIDR®, Zoetis, Parsippany, NJ, USA) for 12–14 d. On day −1 (the day before SCT for fresh transfers), CIDRs were removed and ewes injected with 400 IU Pregnecol® (Bayer, Germany) to induce ovulation. Embryos were transferred to recipient ewes in groups of 2–4 in minimal volume of Embryo Hold medium using a sterile Tomcat catheter and 1 mL syringe. Pregnancies were monitored using ultrasound throughout gestation with the M-turbo ultrasound system (SonoSite, Bothell, WA, USA). Establishment of pregnancy was determined using transrectal ultrasound (L52x, 10-5MHz transducer) between day 30–45 of gestation. Scans after day 45 were completed using transabdominal ultrasound (C60x, 5-2MHz transducer). Scanning was completed approximately once a month, with more frequent scanning if pregnancies were experiencing issues (such as hydrops). Pregnancies were manged by veterinarian and farm staff, where the health of the ewe and the lamb were closely monitored.

Birth was induced hormonally for delivery on day of gestation (D) 147–148. The induction protocol was optimised based on international best practice [24]. The original induction protocol involved 20 mg short-acting dexamethasone phosphate (Dex) administered twice on D147/D148 to initiate parturition. The final protocol used a combination of 5 mg long-acting dexamethasone trimethylacetate (TMA) on D138 for lung maturation, 6 mg Dex on D144 to prime ewes for final Dex dose, and two doses of 20 mg Dex on D145 to initiate parturition. Iterations between original and final protocols varied in presence of priming dose and volume of final Dex doses. Delivery was non-assisted natural or slaughter caesarean. Lambs were bottle fed as per standard farm practice and kept under close veterinary supervision, due to their increased risk status of suffering cloning-related abnormalities.

### *CMAH*/*GGTA1* mutant genotype and phenotype

#### Sample preparation

Blood was collected from all five OFF4 #56 *CMAH*^-/-^ *GGTA1*^-/-^ ewes (1917, 1918, 1919, 1920, and 1921) for genotype and phenotype analysis. For peripheral blood mononuclear cell (PBMC) isolation, blood was collected in purple-top (EDTA) vacutainers (367844, BD Biosciences, Franklin Lakes, NJ, USA), immediately inverted, and stored at room temperature until isolation performed. Room temperature whole blood (3 mL) was diluted 1:1 with PBS and mixed thoroughly. Diluted blood was carefully layered over 4 mL Histopaque (10771, Sigma-Aldrich) in a 15 mL Falcon tube and centrifuged at 400 x g for 30 min at 20°C with the brake off (deceleration rate 0). The PBMC layer was carefully removed, transferred to a fresh 15 mL tube, and washed with PBS up to 15 mL. Cells were centrifuged at 250 x g for 10 min, with brake on (deceleration rate 9). Following centrifugation, the supernatant was removed and hypotonic red blood cell lysis performed by incubating cells in 9 mL MilliQ H_2_O for less than 1 min, followed by addition of 1 mL 10x PBS and mixed immediately. Cells were centrifuged at 250 x g for 10 min. Supernatant was removed and cells washed a second time in 10 mL PBS and centrifuged. Cells were resuspended in PBS, and 5 x 10^5^ cells were transferred to 1.5 mL Eppendorf tubes for staining or DNA extraction.

Samples from various tissues (skin, muscle, heart, liver, kidney, brain, gonad, and spleen) were collected from deceased lambs, dissected into small fragments, and snap frozen in liquid nitrogen. Samples were stored at -80°C. DNA for genotyping was isolated from snap frozen kidney samples using DNeasy Blood and Tissue kit (QIAGEN) according to the manufacturer’s instructions, including additional steps of RNase A incubation and second final elution.

#### Confirmation of lamb sequence

Genotyping was performed on both living and deceased lambs. DNA was amplified using endpoint PCR for *CMAH* and *GGTA1* and sequenced as previously described. For sequences with two edits that could not be separated on a gel, PCR products were subcloned into bacteria using pGEM®-T Easy Vector Kit according to the manufacturer’s instructions. Bacterial colonies were picked and amplified overnight in 3 mL LB. Plasmids were isolated with the Miniprep kit and sequenced using the M13_f primer supplied by the Massey sequencing facility. Sequences were analysed on Geneious.

#### Antigen staining

PBMCs were analysed from all five OFF4 #56 *CMAH*^-/-^ *GGTA1*^-/-^ ewes, one cloned ewe from a Poll-Dorset fetal fibroblast line (‘PDFF’; 1941), and one ewe from artificial insemination (AI; 208), all born in the same season and grazed together. PDFF and AI samples served as positive controls for both Neu5Gc and α-Gal.

Neu5Gc staining was performed by pelleting PBS washed PBMCs at 1,000 x g for 3 min. Supernatant was removed and cells resuspended in 100 µL Neu5Gc primary antibody (1:500) in 0.5% Neu5Gc blocking solution (Anti-Neu5Gc Antibody Kit, 146901, Biolegend, San Diego, CA, USA). Samples were performed in duplicate and a single tube of control WT cells was resuspended in 100 µL chicken isotype control as a negative control (supplied in the Anti-Neu5Gc Antibody Kit). Cells were incubated on ice at 4°C for 1 h. All remaining centrifugation steps were performed at 1,000 x g for 3 min at 4°C.

Following incubation, cells were pelleted by centrifugation, washed with 1 mL PBS, centrifuged again, and supernatant removed. All cells were resuspended in 100 µL of 1:1000 goat anti chicken 488 secondary antibody (2 mg/mL stock; A11039, ThermoFisher) in 0.5% Neu5Gc blocking solution and incubated for 1 h at 4°C. Cell centrifugation and PBS wash steps were repeated. Finally, cells were resuspended in 200–500 µL 0.5% Neu5Gc blocking solution.

α-Gal staining was performed by pelleting PBS washed cells at 1,000 x g for 3 min. Supernatant was removed and cells resuspended in 500 µL of 6 µL/mL isolectin IB_4_ FITC conjugate in PBS (1 mg/mL stock, ALX-650-001F-MC05, Enzo Life Sciences, Farmingdale, NY, USA). Samples were performed in duplicate and a single tube of control WT cells was resuspended in PBS alone as a negative control. Cells were incubated for 1 h at 4°C. Following incubation, cells were pelleted by centrifugation at 1,000 x g for 3 min at 4°C, washed with 1 mL PBS, centrifuged again, and supernatant removed. Stained cells were resuspended in 200–500 µL PBS.

#### Flow cytometry

Stained cells were analysed on the FACSVerse™ flow cytometer (BD Biosciences). Cell populations were gated based on typical forward scatter (FSC) and side scatter (SSC) of PBMCs to avoid measurement of unwanted other cell types, doublets, and debris. Number of gated events was set at 10,000 and fluorescence gating was set on the upper limit of negative controls and lower limit of positive controls.

### Breeding

To generate a flock of *CMAH*^-/-^ *GGTA1*^-/-^ sheep, the five female ewes underwent two different breeding procedures: 1) natural mating and 2) ovum pick-up (OPU), *in vitro* fertilisation (IVF), and ET. Natural mating was performed by introducing a ram for two consecutive estrus cycles.

OPU was carried out by a commercial provider (Animal Breeding Services NZ). Briefly, DKO and WT ewes were synchronised using CIDRs for 10 d, with administration of 0.5 mL Ovuprost (125 µg; Bayer New Zealand Ltd, NZ) on day 6 after CIDR insertion, 2 mL follicle stimulating hormone (FSH; 32 mg/mL) twice (morning and evening) on day 7 after CIDR insertion, and 1 mL FSH twice on day 8 after CIDR insertion. Oocytes were aspirated from follicles using laparoscopy on day of CIDR removal, washed and filtered in IVM aspiration medium, and matured *in vitro* using the same procedure as for abattoir-derived oocytes [25].

After 24–26 h, matured oocytes were processed for IVF. Culture media was prepared by supplementing 8 mL base IVF medium (107.7 mM sodium chloride, 7.15 mM potassium chloride, 0.3 mM monopotassium phosphate, 25 mM sodium bicarbonate, 0.33 mM sodium pyruvate, 3.32 mM sodium lactate, 1.71 mM calcium chloride dihydrate, 8 mg/mL fatty-acid free bovine serum albumin [BSA; MP Biologicals, Auckland, NZ], and 0.04 mM kanamycin sulphate) with 2 mL heat inactivated estrus sheep serum (Day 1.5 post estrus), 0.2 mM penicillamine, and 0.1 mM hypotaurine. Frozen sperm were warmed for 30 s in a 33–37°C water bath, layered on a 40/80% Bovipure gradient, and centrifuged at 300 x g for 15 min. The sperm pellet was then carefully moved to a new tube, washed gently with room temperature Hepes-buffered SOF (HSOF), and centrifuged at 300 x g for 5 min. Supernatant was removed and sperm pellet resuspended in warm IVF medium. Oocytes were washed twice through HSOF, lightly stripping cumulus cells to expose at least 40% of the zona pellucida. Five oocytes were transferred to IVF medium drops with a final volume of 40 µL. Finally, 10 µL sperm were added to each drop for a final concentration of 2 x 10^6^ sperm/mL. Oocytes were incubated at 38.5°C in a modular incubation chamber containing a 5% CO_2_, 7% O_2_, 88% N_2_ gas mixture.

After 18–22 h, zygotes were processed for IVC. Cumulus cells were removed from zygotes using 0.5% pronase (Sigma-Aldrich) in HSOF with 1 mg/mL polyvinyl alcohol, followed by two HSOF washes. Zygotes were then transferred to ESOF plates and cultured as per SCT *in vitro* embryo culture [25]. Blastocysts were morphologically graded on day 6 or 7 before ET, as above [23].

### Statistics

The mean was determined for each indel in the TIDE analysis from pooled transfections in OFF3 and OFF4 (*n*=5; 1x *CMAH*/*GGTA1* OFF3, 1x *CMAH*/*GGTA1* OFF4, 3x *CMAH*/*GGTA1*/another unrelated gene OFF3). Error bars represent standard error of the mean (SEM). Overall efficiency is displayed as the mean for the five individual efficiencies. R^2^ value (fit of prediction to data determined by TIDE) shown as the lowest value for all transfections, indicating the worst predicted fit of the five samples.

Reconstructs generated across SCT runs within a year were pooled. The number of cleaved embryos was normalised to the number of reconstructs placed into *in vitro* culture (IVC). Total blastocyst development rate was normalised on cleaved embryos. *P* values for SCT and ET were calculated using Fisher’s exact test in Excel. Flow cytometry results were presented as the percentage of events counted within the fluorescent positive gating out of total counted within PBMC gating. The mean was calculated for the samples run in duplicate ± the standard deviation.

## Results

### gRNA-Cas9-mediated disruption of *CMAH* and *GGTA1*

Since sheep genetic variation is not well annotated in public datasets, target sequences for *CMAH* (Fig. S1a) and *GGTA1* (Fig. S1b) were confirmed in both ovine fetal fibroblast cell lines (OFF3 and OFF4) to inform gRNA design. Three *CMAH* gRNAs were designed to cut within 5 bp of each other, while a single gRNA was used for targeting *GGTA1* after previously being identified as the most efficient in goat (C. Di Genova, unpublished). All guides were ranked by off- and on-target activity (Table S1a) and editing efficiency determined using PCR. Cas9 was used to introduce non-homologous end joining mutations in exon 5 of *CMAH* (Fig. 1a) and the final exon of *GGTA1* (Fig. 1b) in OFF3 and OFF4. Editing efficiency of the three *CMAH* gRNAs in OFF3 was determined as 94.6%, 94.3%, and 88.1% for gRNA1, 2, and 3, respectively (Fig. S2). While no WT sequences were detected from the sample edited with gRNA1, low levels were detected following transfection with both gRNA2 (1.1%) and gRNA3 (4.0%). As gRNA1 had the lowest WT detected, it was used to target *CMAH*. Following transfections into OFF3 and OFF4, the average editing efficiency with *CMAH* gRNA1 was 90 ± 1% (R^2^≥0.89, *n*=5, Fig. 2a). No WT sequences were detected in any of the transfections. The most common edits were deletions <5 bp and a single base insertion. *GGTA1* editing efficiency was 93 ± 1% across all transfections in OFF3 and OFF4 (R^2^ ≥0.92, *n*=5) with no WT sequences detected (Fig. 2b). The most common edit was a single base insertion (>50% edits detected) and most deletions were <5 bp, except for a 20 bp deletion in 11% of the initial OFF3 transfection.

**Figure 1.**
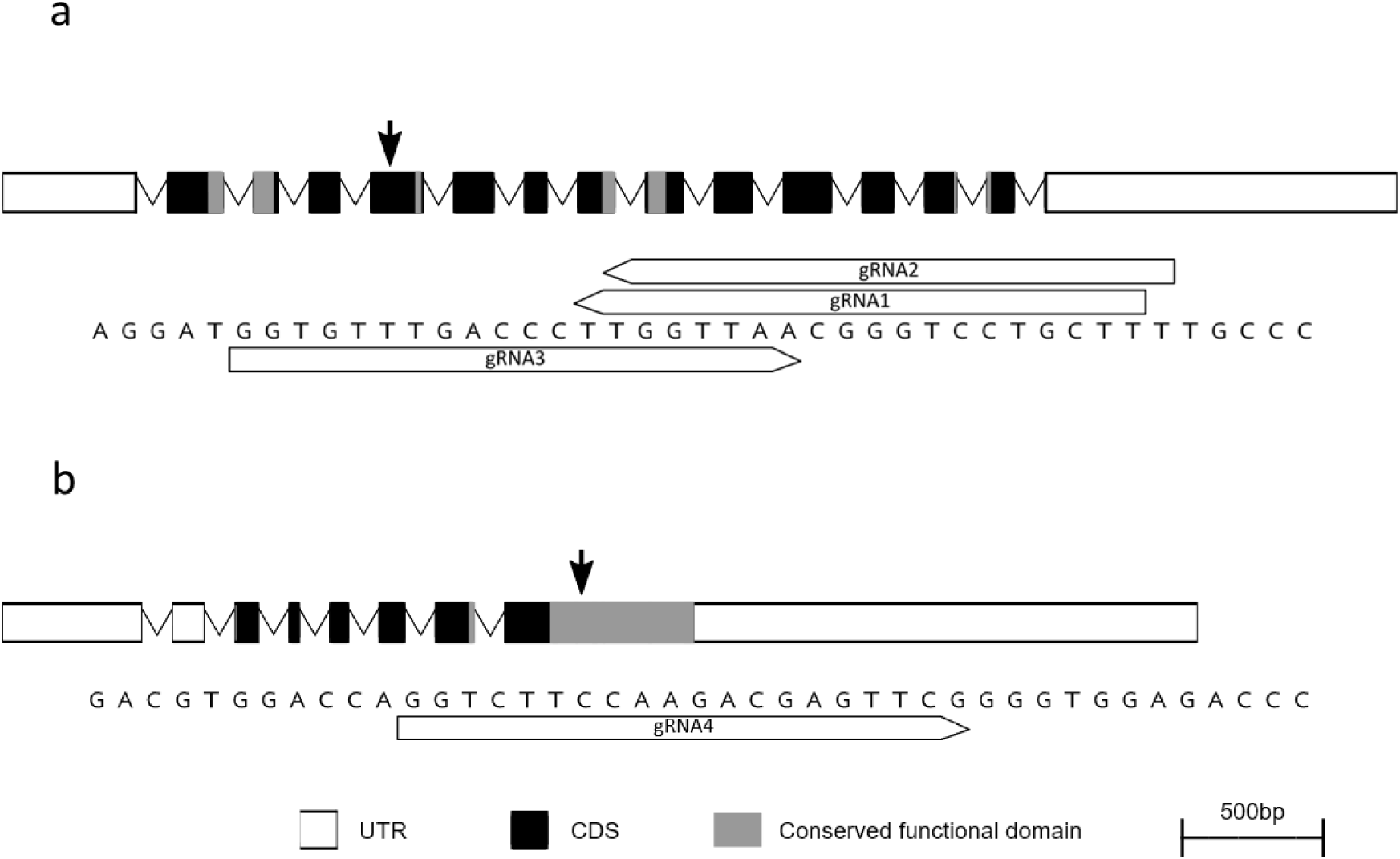
Ovine gene structure and editing locations for (a) *CMAH* and (b) *GGTA1*. Gene structure shows introns (lines) and exons (boxes) containing coding sequence (CDS, black) and conserved functional domains (grey). Previous target sites from the literature indicated by grey arrows. Target site for Cas9 editing indicated by black arrow and sequence displayed below gene structure, along with gRNA binding sites.

**Figure 2.**
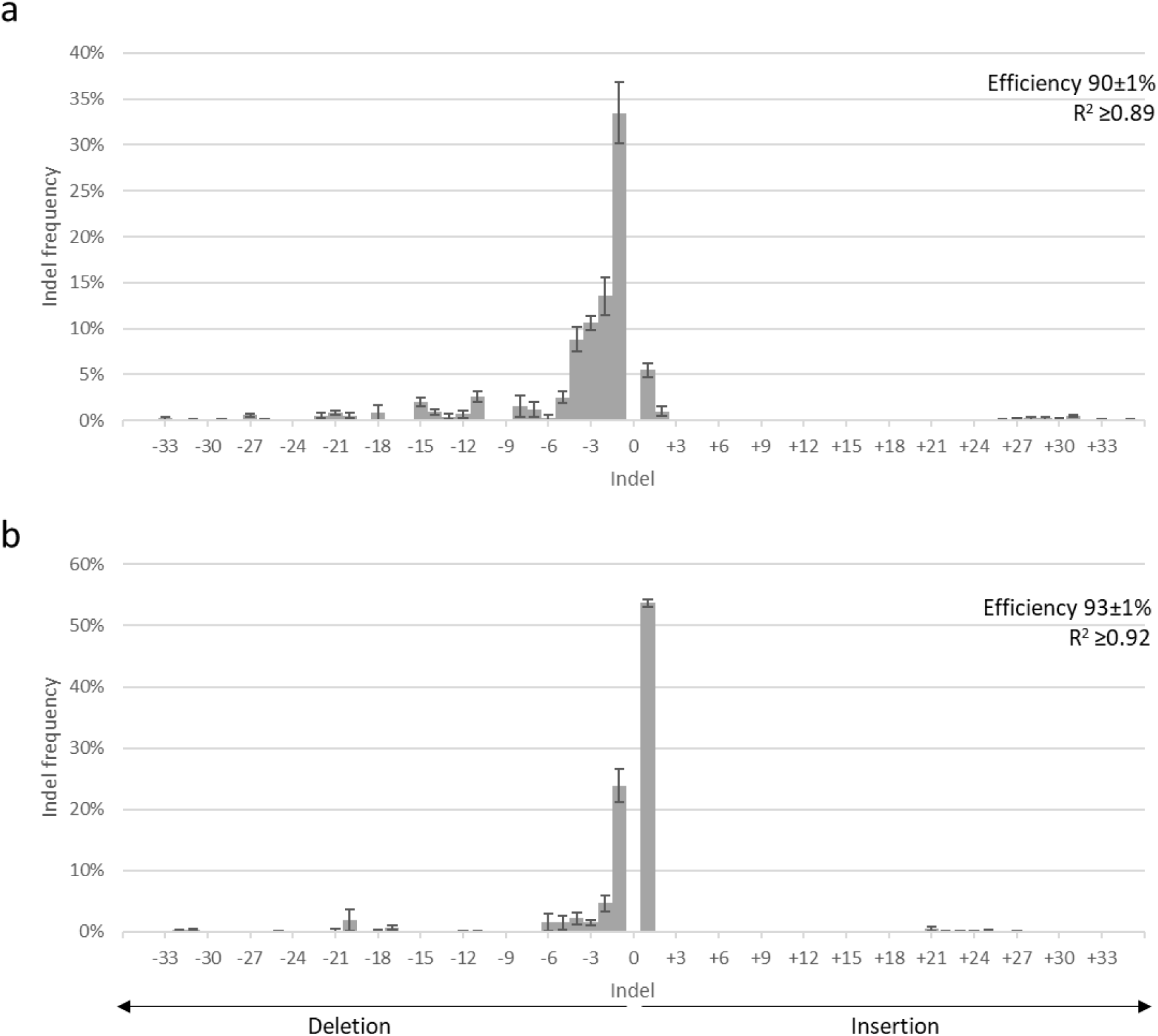
TIDE analysis of editing in OFF3 and OFF4 for (a) *CMAH* gRNA1 and (b) *GGTA1*. Sanger sequencing results of edited population from each transfection was run through TIDE to estimate the proportion of indels present. Error bars = SEM, n= 5.

DKO clonal cell strains were isolated using manual selection of mitotic doublets and screened via Sanger sequencing for mutations in both genes. Of the 107 OFF3 mitotic cells selected, 77 (72%) grew to confluency in 96-well plate and 61 (57%) proliferated after passaging (Table S2a). Of these strains, 55 were sequenced; no band was observed for *CMAH* in 6 strains, which were excluded from further analysis. Isolation of OFF4 strains involved selection of 90 mitotic cells, with 83 (92%) growing to confluency. At this point 61 (68%) strains were contaminated and removed from culture. Of the remaining strains, 16 (55%, 16/29 non-contaminated strains) proliferated after passaging. Sanger sequencing revealed 11/12 mutation profiles were clonal in the OFF3 strains and 10/14 in OFF4 (Table S2b). Three compound heterozygous DKO strains (#11, #66, and #56) were chosen for SCT cloning.

A representative strain of each mutation profile was assessed for integration of the Cas9 plasmid sequence. A positive PCR band was detected for 1/10 OFF3 (Fig. S3a) and 5/14 OFF4 strains (Fig. S3b) and four OFF3 strains had faint bands, meaning Cas9 integration could not be excluded. Of the strains conclusively not containing Cas9 sequence, four OFF3 strains and a single OFF4 strain were screened for off-target Cas9 effects. No mutations were detected via Sanger sequencing of the top three off-target sites determined by CRISPOR (Fig. S3c,d). Two off-target sites (*CMAH* site 2 and *GGTA1* site 3) contained SNPs in the OFF4 WT sequence compared with OFF3 WT and the sheep reference genome, which were maintained in strain #56.

### Generation of *CMAH/GGTA1*-deficient lambs

Cleaved SCT reconstructs developed into blastocysts at similar rates across DKO strains within each year (2018 #11 75/395=19%, #66 34/321=15%, *p*=0.21; 2019 #66 51/621=8%, #56 46/501=9%, *p*=0.64; Table 1). In 2018, strain #66 non-aggregated morula developed into high grade blastocysts more frequently than #11 (23/34=68% vs 22/75=29%; *p*=0.0004).

**Table 1.** Somatic cell cloning development of OFF3 (#11/#66) and OFF4 (#56) *CMAH*^-/-^ *GGTA1*^-/-^strains.

| Strain | Year | n<sup>1</sup> | nIVC<sup>2</sup> | >1-cell<br>(%)<sup>3</sup> | Non-aggregate<br>blastocysts | Aggregate<br>blastocysts | B 1–3 (%)<sup>4</sup> | B 1–2 (%)<sup>5</sup> |
| --- | --- | --- | --- | --- | --- | --- | --- | --- |
| #11 | 2018 | 12<sup>A</sup> | 530 | 395 (75) | 75 | n/a | 75 (19) | 22 (29) |
| #66 | 2018 | 6 | 293 | 231 (79) | 34 | n/a | 34 (15) | 23 (68) |
|  | 2019 | 4<sup>B</sup> | 735 | 621 (84) | 8 | 43 | 51 (8) | 24 (47) |
| #56 | 2019 | 4<sup>B</sup> | 626 | 501 (80) | 8 | 32 | 46 (9) | 21 (46) |
| **Total** |  | **26** | **2184** | **1748 (80)** | **125** | **75** | **206 (12)** | **90 (44)** |

<sup>1</sup> Number of somatic cell cloning runs.
<sup>2</sup> Number of embryos placed into *in vitro* culture (IVC).
<sup>3</sup> Cleavage normalised on nIVC.
<sup>4</sup> Total blastocyst development normalised on > 1-cell embryos.
<sup>5</sup> Proportion of grade 1–2 blastocysts normalised on total number of blastocysts.
<sup>A</sup> Three runs used *in vivo* matured oocytes.
<sup>B</sup> Embryo aggregation at morula stage was performed in 2019 runs.

From the two male DKO strains #11 and #66, 1/50 (2%) and 5/78 (6%) transferred blastocysts generated lambs, respectively (Table 2). However, no male lambs survived to weaning. The female #56 strain was more successful, with 8/40 (20%) transferred blastocysts generating lambs (*p*=0.012 vs #11, *p*=0.060 vs #66), with 5 lambs surviving. The majority of cloned lambs died perinatally (7/14=50%) or were euthanised on welfare grounds (2/14=14%) (Table S3). Deceased lambs displayed a range of abnormalities in various organ systems (musculoskeletal, pulmonary, hepatic, renal, intestinal and cardiovascular), which are typical cloning-associated phenotypes (Table S4).

**Table 2.** ***In vivo* survival of *CMAH*^-/-^ *GGTA1*^-/-^ cloned embryos.**

| Cell line | Recipient ewes | Embryos<br>transferred | Lambs born (%)<sup>1</sup> | Lambs alive at<br>weaning (%)<sup>1</sup> |
| --- | --- | --- | --- | --- |
| OFF3 #11 | 17 | 50 | 1 (2%) | 0 (0%) |
| OFF3 #66 | 28 | 78 | 5 (6%) | 0 (0%) |
| OFF4 #56 | 15 | 40 | 8 (20%) | 5 (13%) |
| **Total** | **60** | **168** | **14 (8%)** | **5 (3%)** |

<sup>1</sup> Normalised on blastocysts transferred.

### *CMAH*/*GGTA1* mutant genotype

Five female lambs survived into adulthood (Fig. 3a). All surviving lambs derived from the OFF4 #56 cell strain and were confirmed to carry the desired *CMAH*/*GGTA1* edits using Sanger sequencing (Fig. 3b). DNA was isolated from snap frozen kidney samples of deceased lambs or PBMCs from live lambs to compare genotypes to the original edited cells. Chromatograms from all lambs matched that of the parental strains edited for *CMAH* (Fig. S4) and *GGTA1* (Fig. S5). A representative from each cell line was sub-cloned into bacteria to identify the exact position of the edit identified by TIDE. All edits matched the size identified by TIDE and confirmed the bases of the single nucleotide insertions. All edits caused frameshift mutations, mostly introducing a premature stop codon shortly after the mutation site. The exception was the single base deletion in *GGTA1* (Allele 1 of OFF3 #11 and OFF4 #56). While it also caused a frameshift mutation, there was predicted 76 amino acids before introduction of a stop codon.

**Figure 3.**
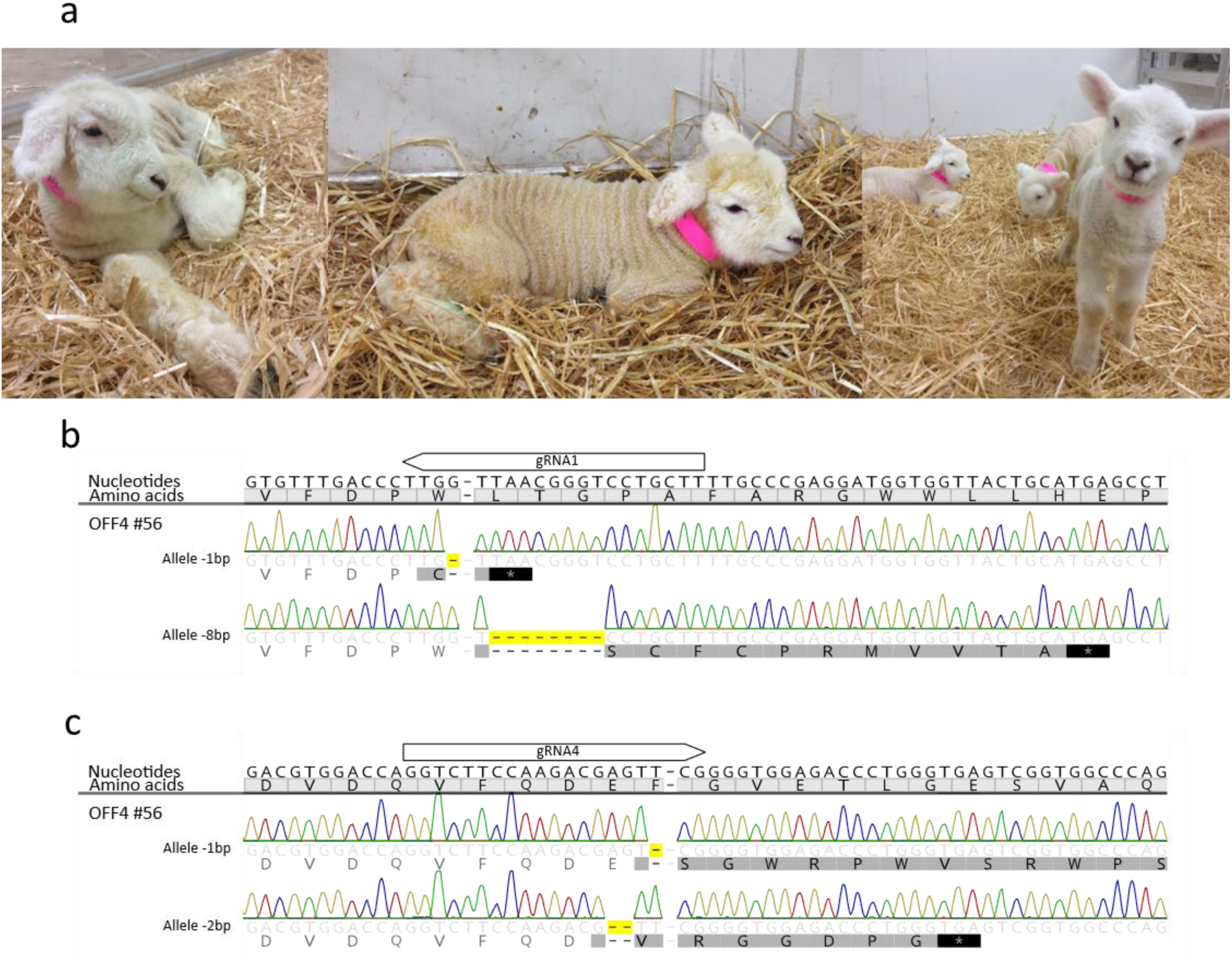
Cloned double knockout (*CMAH^-/-^ GGTA1^-/-^*) lambs and their genotype. All lambs were derived from OFF4 #56 cell strain. (a) Lambs at <1 week old. From left to right 1917, 1918, 1921, 1920, and 1919 (foreground). Sanger sequence of cloned lambs for (b) *CMAH* and (c) *GGTA1*. Translation is shown under the nucleotide sequence. Nucleotides and amino acids that differed from the wild-type are shown in black font, matching nucleotides and amino acids are shows in light grey font. Yellow boxes highlight edited nucleotides. Asterisks indicate stop codons.

### *CMAH*/*GGTA1* mutant phenotype

PBMC samples from the five living ewes were analysed for absence of Neu5Gc and α-Gal. Control samples were collected from a PDFF cloned ewe (1941), that underwent the same SCT cloning, ET, and rearing procedures as the edited sheep, and an AI ewe (208) that was born within the same season and grazed alongside the cloned sheep. All five edited ewes were negative for Neu5Gc primary antibody (positive PBMCs: 1917 = 1.1 ± 0.1%, 1918 = 1.1 ± 0.1%, 1919 = 0.6 ± 0.1%, 1920 = 0.5 ± 0.1%, 1921 = 0.7 ± 0.1%; *n*=2) compared with their WT counterparts (1941 = 97.4 ± 2.3%, 208 = 99.8 ± 0.1%; *n*=2) (Fig. 4a). Edited sheep showed a similar profile to the negative control (AI ewe treated with the chicken isotype primary antibody control; 1.6%, *n*=1). Edited ewes were also negative for α-Gal (positive PBMCs: 1917 = 2.5 ± 0.2%, 1918 = 2.4 ± 0.7%, 1919 = 2.6 ± 0.2%, 1920 = 2.1 ± 0.7%, 1921 = 2.1 ± 0.7%; *n*=2) compared with WT controls (1941 = 70.5 ± 1.1%, 208 = 82.2 ± 0.6%; *n*=2) (Fig. 4b).

**Figure 4.**
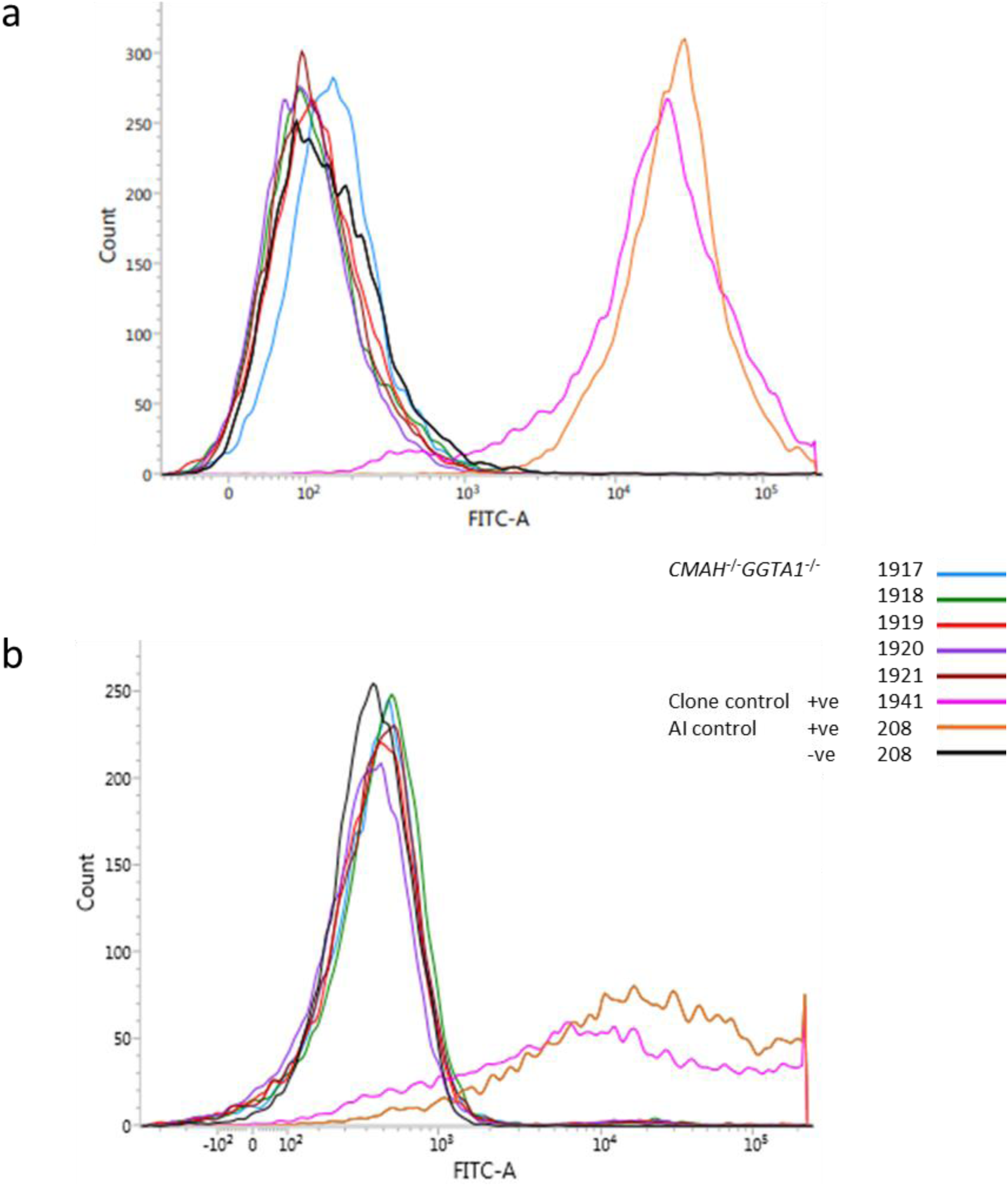
Confirmation of (a) Neu5Gc and (b) α-Gal absence in the five OFF4 #56 *CMAH^-/-^ GGTA1^-/-^* ewes. PBMCs stained with Neu5Gc antibody or isolectin B4 FITC conjugate, respectively. Positive control (+ve) samples from a PDFF cloned ewe (1941) and an AI ewe (208). Negative controls (-ve) were AI 208 incubated with chicken isotype control for Neu5Gc or no isolectin for α-Gal.

Edited sheep were similar to the negative control (AI 208 PBMCs that underwent all washing and incubation steps without addition of isolectin; 0.4%, *n*=1).

### Breeding *CMAH*/*GGTA1* mutant ewes

To generate a *CMAH*^-/-^ *GGTA1*^-/-^ DKO flock, the five OFF4 #56 ewes underwent two rounds of natural mating with WT rams. In both years, no DKO ewes became pregnant after running them with a ram for two consecutive estrus cycles. By contrast, all WT controls (n=3) became pregnant and lambed without incident (Table S5).

To evaluate for potential fertility issues in the cloned DKO ewes, OPU/IVF/ET was performed. Oviducts and uteri were externalised during laparotomy and appeared anatomically normal upon visual inspection by experienced veterinarians carrying out the OPU procedure. OPU was performed twice on each DKO ewe, with an average of 12.6 follicles aspirated and 12 oocytes recovered per donor (Table S6). This recovery rate was similar to the WT ewes (11.9 follicles per donor and 10.9 oocytes per donor). IVF results were similar for DKO and WT oocytes (Table S7) with 22% and 31% of cleaved embryos developing into blastocysts (*p*=0.402) from DKO and WT oocytes, respectively, comprising 60% and 75% high-quality blastocysts (*p*=0.705). Following ET of these blastocysts into WT surrogates, 2 lambs were born from DKO-derived oocytes (Table S8). Unfortunately, both lambs were euthanised on welfare grounds. Development to term was similar for embryos from DKO oocytes compared with WT oocytes (2/9=22% vs 11/36=31%; *p*=0.964), with no large differences in lamb survival for DKO oocytes (0/9=0% vs 7/36=19%; *p*=0.368). We conclude that DKO ewes displayed normal ovarian function and produced fully developmentally competent oocytes, indicating that their inability to establish viable pregnancies after AI is due to extrinsic factors, not to quality of the heterozygous *CMAH/GGTA1* embryo.

## Discussion

This is the first demonstration of a *CMAH/GGTA1* DKO phenotype in a live sheep model. We confirm that *CMAH* and *GGTA1* are necessary to form the Neu5Gc and α-Gal antigens, respectively, with no functional compensation by other genes when these enzymes are absent. Compared to pig, sheep offer advantages in terms of improved IVP efficiency [26] and lower embryo transfer requirements [27], as well as wider cross-cultural acceptance. As discussed below, this refines sheep as an alternative model for clinical transplant testing and xenotransplantation.

### Phenotype of edited lambs

Both male OFF3-derived cell strains failed to produce viable animals while the female OFF4 strain was successful. In contemporaneous experiments, other cell strains from the OFF3 parental cell line produced healthy *NANOS2*^-/-^ mutant lambs [28], indicating no inherent genetic problems with this line. Pathologies in the deceased male cloned lambs were consistent with the cloning syndrome, including defects in the pulmonary, renal, musculoskeletal, hepatic, and cardiovascular systems [29,30]. Therefore, this was likely a generic cloning-related problem, rather than a specific artefact related to *CMAH* and/or *GGTA1* editing.

The loss of all male DKO lambs was a setback for producing a flock of xenocompatible sheep, requiring the generation of an intermediate, heterozygous F1 generation from the DKO ewes. This further complicates breeding to ensure homozygous transmission of both independently segregating KO alleles. As a first step, we mated DKO ewes with different WT rams of proven fertility to generate heterozygous *CMAH*^+/-^ *GGTA1*^+/-^ embryos *in vivo*. All ewes, including WT controls, showed normal signs of heat and were repeatedly serviced by the ram, as indicated by their colour markings, over two consecutive estrus cycles in alternate mating seasons. However, none of the DKO ewes were detected as pregnant using ultrasonography around one and three months after natural mating with a ram, while all WT surrogates showed fetal heartbeat at scanning and lambed without incident.

Since there were no discernible issues with the rams’ and ewes’ sexual behaviour over the mating period, we focused on potential fertility issues in the cloned DKO ewes, either in their reproductive tracts or gonads. There was no indication of ovarian malfunction and DKO oocytes were fully competent to develop into blastocysts after IVF with WT sperm. Likewise, DKO-derived IVF embryos developed into live offspring similar to WT controls. Thus, heterozygous *CMAH*/*GGTA1* embryo quality and development appeared normal in a WT reproductive environment.

By contrast, none of these embryo genotypes survived in the uterus of DKO surrogates. We hypothesize that this is related to the presence of antibodies in the cloned DKO ewes that target α-Gal and Neu5Gc xenoantigens present on the embryo surface. GalKO pigs [31] and sheep [15] express anti-Gal antibody at clinically relevant levels. By analogy, it is likely that all adult DKO ewes produced both cytotoxic anti-Gal and anti-Neu5Gc antibodies. The α-Gal epitope is uniformly present on the surface of trophoblast cells in mammalian blastocysts where it is recognised by the antibodies in GalKO pig serum, leading to rapid (within minutes) complement activation and cell lysis in immunosurgery assays [31]. It is unclear to what extent antibody-dependent cell cytotoxicity may already compromise blastocyst viability before implantation when there is no direct contact with maternal blood, but immune surveillance occurs through the local mucosa and uterine endometrial cells to mediate maternal recognition of pregnancy [32]. Around implantation, maternal blood perfusion begins and concomitant exposure to circulating antibodies increases the risk of rejection, which is consistent with the lack of any observed implantation events upon transferring heterozygous *CMAH/GGTA1* embryos into DKO ewes. These issues only came to the fore due to the lack of DKO males, which prompted us to mate DKO ewes with WT rams. A more typical breeding scenario would be the reciprocal situation, whereby WT females are bred with KO males, followed by crossing the heterozygous female offspring to KO males to obtain homozygous progeny. This avoids any immune complications associated with breeding KO females to WT rams.

### Sheep as a model for xenotransplantation and clinical testing

Xenocompatibility of *CMAH*^-/-^ *GGTA1*^-/-^ sheep cells will need to be assessed in preclinical *in vitro* and *in vivo* trials. This includes flow cytometric analysis of reduced cross-reactivity between ovine DKO vs WT PBMCs with human sera [9] and transplantation of DKO sheep cells into α-Gal negative mice without eliciting an immune response [6]. Lowered immune response in these studies would be followed by organ perfusion with human blood to evaluate the tolerance of human cells to maintain normal function [33]. To reach clinical trials in New Zealand, xenotransplant donors need to have completed preclinical studies in at least two animal models, including one non-human primate [34].

Continued optimisation of sheep cells for transplantation into humans would entail further genetic modifications. Candidate KOs to add include *CD46* [35], *β4GalNT2* [36] and/or *CIITA* [37], as well as human transgene knock-ins [38]. In pig, these multi-gene modifications culminated in landmark xenotransplantation trials, including the first pig-to-human heart transplant using a donor pig with ten gene edits (Revivicor), including four pig gene KOs (*GGTA1*/*CMAH*/*β4GalNT2* and growth hormone receptor [*GHR*]) augmented with six human transgene knock-ins to render the cardiac xenograft more immunocompatible [38]. More recently, transplantation of a porcine kidney with 69 genomic edits, including deletion of *GGTA1*/*CMAH*/*β4GalNT2,* inactivation of porcine endogenous retroviruses, and insertion of seven human transgenes, resulted in sustained kidney function without evident xenograft rejection, albeit under an expanded immunosuppression regime [39]. These models now define the gold standard for clinical transplants, achieving unprecedented survival times in living patients.

Both sheep and pigs have comparable size and physiology to humans, making them ideal donor species. Sheep and pigs also share similar growth rate and short pregnancies (four vs five months for pigs and sheep, respectively) resulting in fast generation of animals. Overall animal generation would be faster with pigs, as they have a high number of offspring (5–12) compared with sheep (1–2). However, ease of *in vitro* production of sheep embryos and a ten-fold decrease in embryos required for transfer (1–4 in sheep vs >50 in pigs), enhances the experimental operation with sheep.

The ethical principle of autonomy dictates that physicians must inform patients from any religion about the source of the organ to be transplanted [40]. So far, pigs have been the most common source of organs for use in human therapeutics, due to similarities in physiology and diet [11]. However, the use of pigs raises ethical, cultural, and religions concerns that need to be addressed. First, many Islamic cultures prohibit transplantation of porcine organs to replace defective human ones [41]. Whilst some Islamic scholars argue that organ transplants may be permissive if it was a life-saving measure [42], an alternative donor species would mitigate this concern. A similar argument would apply to Jewish patients, even though they can more freely use porcine products as live-saving organ transplants [43]. Both religious groups may prefer sheep organs, if such a choice was available.

Another concern with using livestock organs for transplantation is the transfer of zoonotic pathogens. Biosecure breeding and housing allows animals to be raised free of microorganisms that could be transferred to humans [11]. Retroviral sequences, integrated into the livestock genome, are an exception. These inactive endogenous retroviruses (ERVs) have been found in all mammals, including sheep [44] and pigs, causing concern of cross-species reactivation when transplanted into humans [45]. There are no reports that ERVs can cause disease in humans and transplant recipients have not shown evidence of infection [38,46]. Even though the threat of transmission to humans is hypothetical, Cas9 editing has been used to KO all copies of pig ERVs in a cell line by targeting a conserved sequence [47]. Similar modifications may be needed to inactivate sheep ERVs before xenotransplantation.

In addition to potential transplant donors, sheep are an essential recipient model for functional testing of durability and regeneration of bioprosthetic and tissue-engineered heart valves, respectively. However, their lack of anti-Gal antibodies has been a major limitation, potentially underestimating the risks associated with immune rejection, inflammation, and premature valve transplant degeneration [48–50]. Since anti-Neu5Gc antibody further increases tissue calcification in a subcutaneous implant model [51], elimination of Neu5Gc-modified glycans from heart valve tissue will likely further reduce immune injury from human antibody binding. The orthotopic sheep implantation model provides reproducible preclinical outcomes that are accepted by regulatory bodies as industry standard. This confers advantages over KO mice, rabbits, or pigs that also produce xenogeneic antibodies and could mimic the immune response of human patients to commercial clinical transplants. Thus, the DKO sheep presented here provide a refined model for the rigorous testing of heart valves, as well as other biomedical devices, such as cochlear implants [52].

Last, sheep-derived products are also a major source of proteins for human consumption and source of dairy products. Immune-compatible sheep may be beneficial for human consumption, reducing α-Gal-associated meat allergies [53], inflammation, arthritis, and cancer [4]. The health benefits of hypoallergenic sheep can add value to future food production with regulatory approval already being granted to *GGTA1* KO pigs by the US Food and Drug Administration [54].

In summary, low immunogenic DKO sheep-derived products can now be further explored for clinical purposes related to xenotransplantation, as well as human consumption.

## Acknowledgments

We thank Axel Heiser and Olivia Wallace for help with flow cytometry and PBMC isolation. Aaron Malthus and Tim Hale took care of animal husbandry, while Elyssa Barnaby and Ali Cullum provided veterinarian care. Ovary collection was performed by Murray Brown, Angela Brennan, Tony Kerema, Katherine Moors, and Sue Odom. Commercial ovum pick-up and embryo transfer was performed by Animal Breeding Services Ltd. This work was funded by a PhD fellowship from the University of Auckland to SJA, with further support by the Todd Foundation and Maurice & Phyllis Paykel.

## Author Contributions

SJA and BO conceived and designed the experiments. SJA carried out most of the experiments, processed and analysed the data, and produced the figures. CDG and SW designed and tested GGTA1 gRNA and Cas9 primers. ZLM derived the parental cell lines. LMF, ZLM, JW, FM and DW contributed to sheep cloning and cryopreservation. SD took care of embryo transfers and animal husbandry. SJA and BO wrote the manuscript. All authors read, provided comments, and approved the final manuscript.

## Competing Interest Statement

The authors declare no competing interests.

## Notes

### Competing Interest Statement

The authors have declared no competing interest.

