## Supplemental Figures and Tables for "Generation of cloned sheep lacking galactose-α1,3-galactose and N-glycolylneuraminic acid antigens"

**This file includes:**

Supplementary Figures 1–5 (Page 2–6)

Supplementary Tables 1–8 (Page 7–15)

### Supplementary Figures

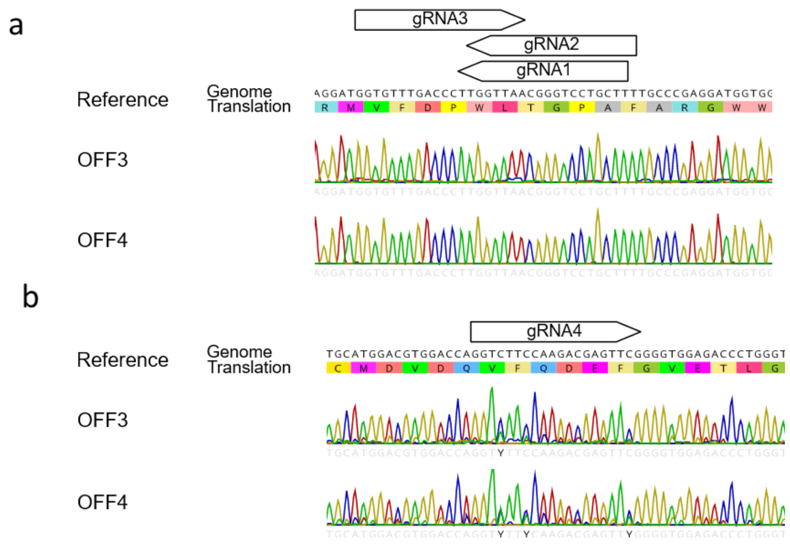


**Figure S1. Confirmation of parental cell line sequences for (a) *CMAH* and (b) *GGTA1*.** Grey shading indicates functional domain in *GGTA1*. Ambiguities in *GGTA1* sequence ("Y") for OFF3 resolved in sequencing with reverse primer.

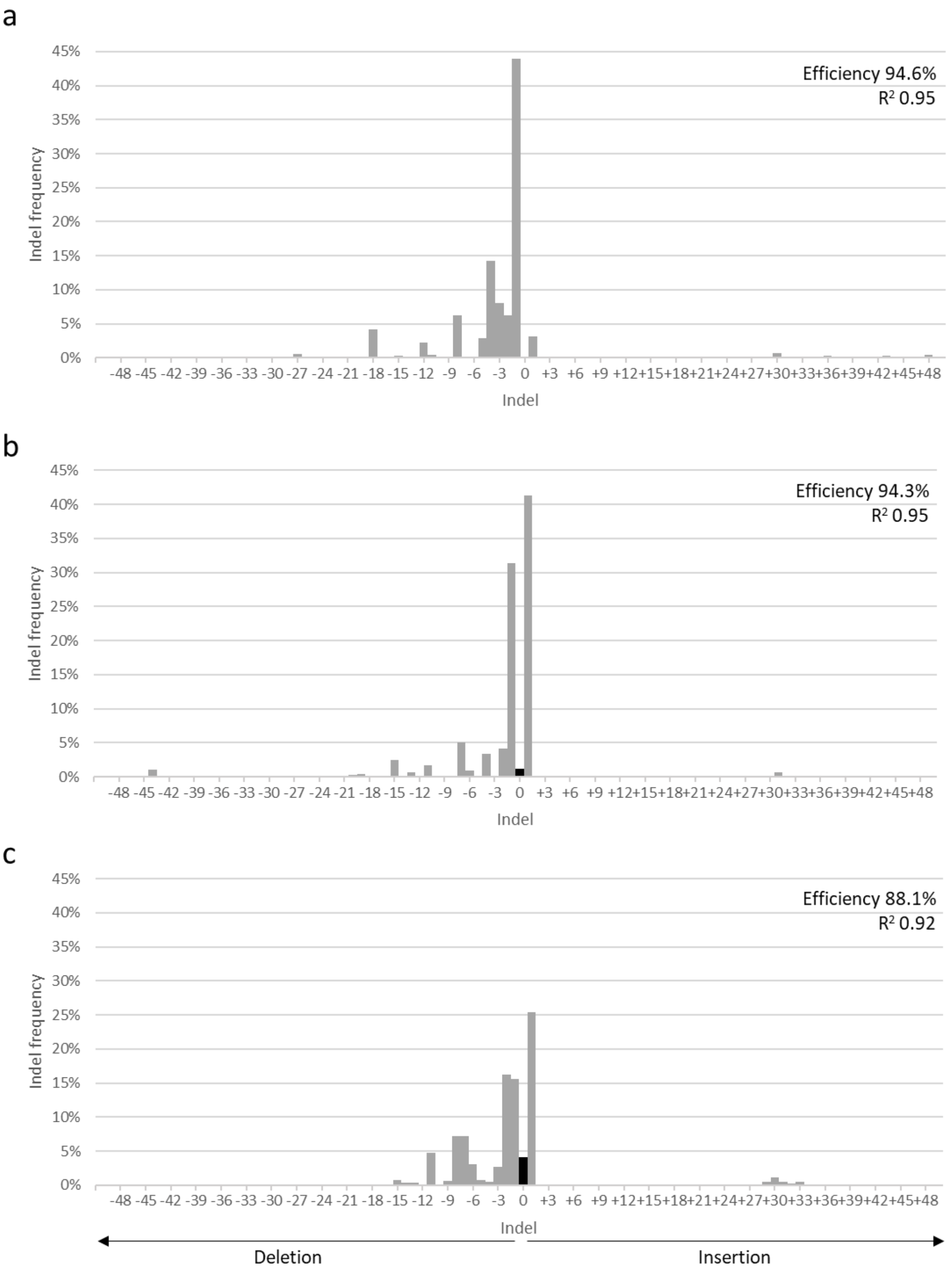


**Figure S2. TIDE analysis of CMAH (a) gRNA1, (b) gRNA2, and (c) gRNA3 from single OFF3 transfection.** Sanger sequencing results of edited population from each transfection was run through TIDE to estimate the proportion of indels present. Unedited sequences shown in black. n= 1.

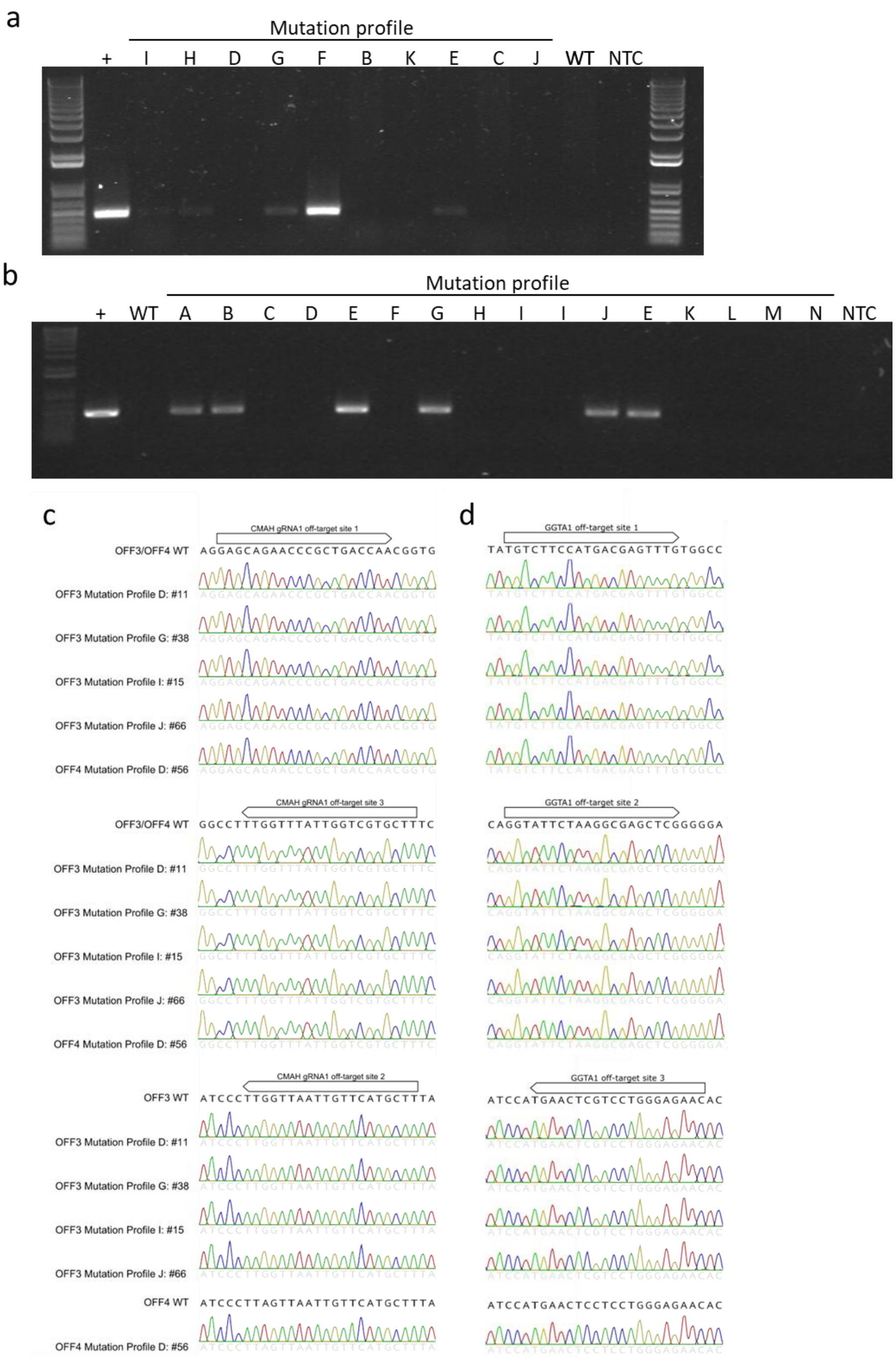


**Figure S3. Cas9 plasmid insertion and off-target screening.** Cas9 plasmid insertion PCR for (a) OFF3 and (b) OFF4 strains. End-point PCR performed with primers designed to Cas9 sequence in plasmid backbone. Positive control (+) PX459. Negative controls wild-type DNA (WT) and no template control (NTC). Off-target screen of top three sites for (c) CMAH gRNA1 and (d) GGTA1 gRNA4 in selected strains. Sanger sequencing of regions identified by CRISPOR as most likely off-target sites. Boxes indicate off-target sequence (not including PAM site). CMAH off-target site 2 and GGTA1 off-target site 3 have a SNP in the WT sequence of OFF3 vs OFF4 (underlined).

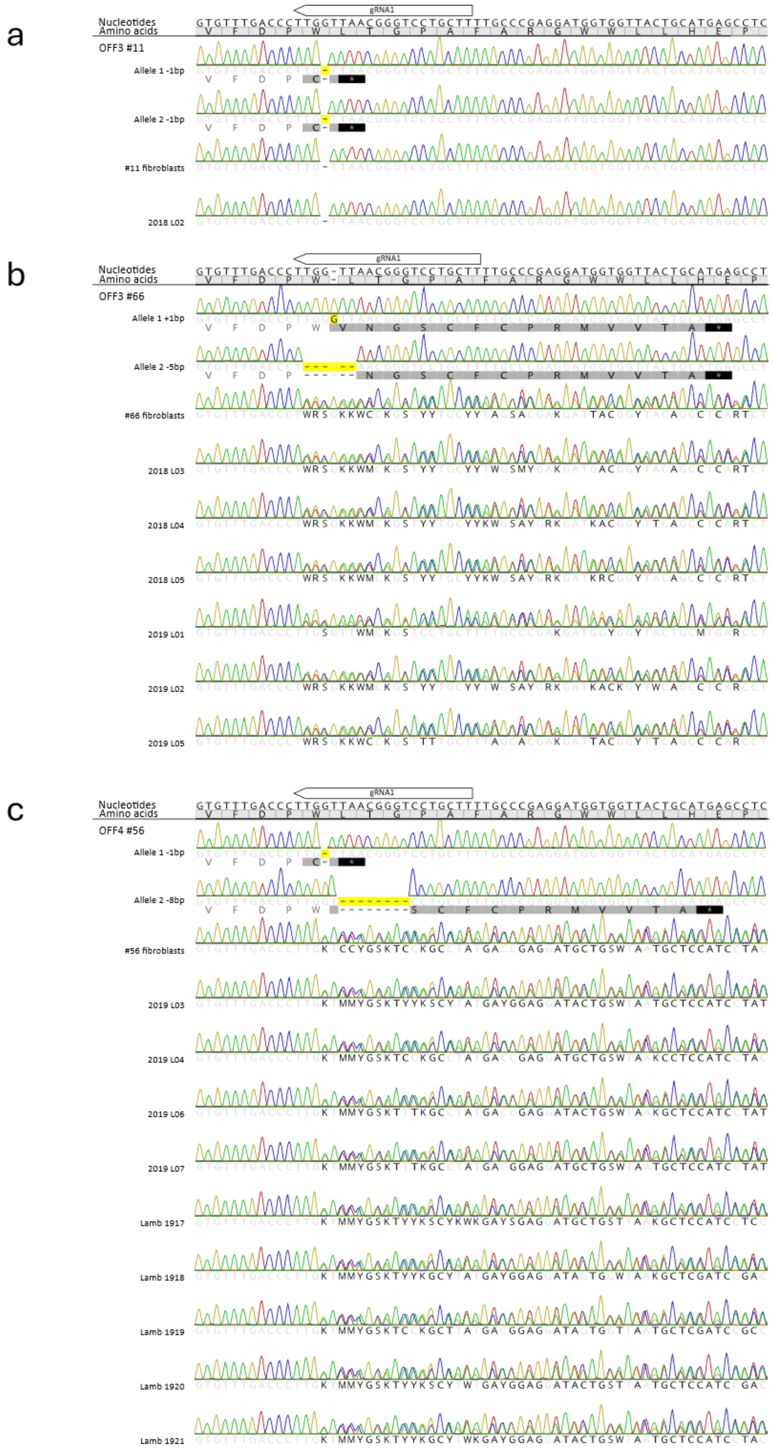


**Figure S4. Sanger sequence of *CMAH* in lambs derived from (a) OFF3 #11, (b) OFF3 #66, and (c) OFF4 #56 cells.** From top to bottom, each panel shows the nucleotide and amino acid sequence of the reference genome and individual edited alleles in each PCR-subclone, followed by sequence reads for each edited fibroblast strain and for each cloned lamb. Nucleotides and amino acids that differed from the wild-type are shown in black font, matching nucleotides and amino acids are shows in light grey font. Yellow boxes highlight edited nucleotides. Asterisks indicate stop codons.

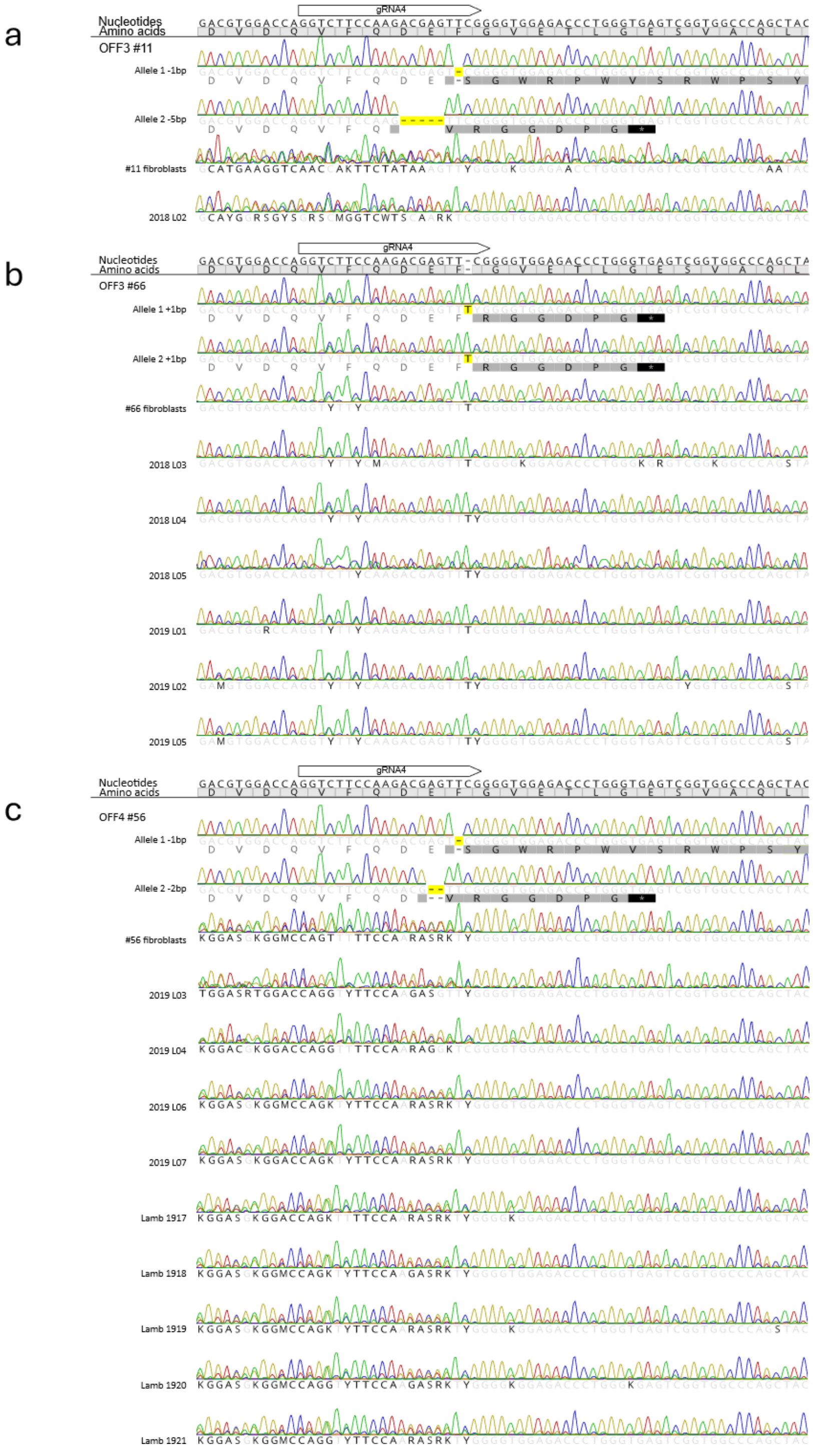


**Figure S5. Sanger sequence of *GGTA1* in lambs derived from (a) OFF3 #11, (b) OFF3 #66, and (c) OFF4 #56 cells.** From top to bottom, each panel shows the nucleotide and amino acid sequence of the reference genome and individual edited alleles in each PCR-subclone, followed by sequence reads for each edited fibroblast strain and for each cloned lamb. Nucleotides and amino acids that differed from the wild-type are shown in black font, matching nucleotides and amino acids are shows in light grey font. Yellow boxes highlight edited nucleotides. Asterisks indicate stop codons.

### Supplementary Tables

**Table S1. (a) CMAH and GGTA1 gRNA sequences and efficiency scores, and (b) gRNA oligo and primer sequences.**

a

| | | | | Geneious10 | | CRISPOR | | |
|---|---|---|---|---|---|---|---|---|
| **Gene** | **gRNA number** | **Sequence** | **PAM** | **Off-target score** | **Activity score** | **Specificity score** | **Predicted efficiency** | **Predicted off-target sites** |
| CMAH | gRNA1 | AAGCAGGACCCGTTAACCAA | GGG | 89.37% | 61% | 96 | 66 | 50 |
| CMAH | gRNA2 | AAAGCAGGACCCGTTAACCA | AGG | 87.84% | 20% | 95 | 64 | 66 |
| CMAH | gRNA3 | GGTGTTTGACCCTTGGTTAA | CGG | 79.63% | 13% | 88 | 37 | 106 |
| GGTA1 | gRNA4 | GGTCTTCCAAGACGAGTTCG | GGG | 93.63% | 42% | 97 | 67 | 31 |

b

| Name | Target | Direction | Sequence | Amplicon size (bp) |
|---|---|---|---|---|
| **gRNA1** | CMAH | f | caccgAAGCAGGACCCGTTAACCAA | |
| **gRNA1** | CMAH | r | aaacTTGGTTAACGGGTCCTGCTTc | |
| **gRNA2** | CMAH | f | caccgAAAGCAGGACCCGTTAACCA | |
| **gRNA2** | CMAH | r | aaacTGGTTAACGGGTCCTGCTTTc | |
| **gRNA3** | CMAH | f | caccGGTGTTTGACCCTTGGTTAA | |
| **gRNA3** | CMAH | r | aaacTTAACCAAGGGTCAAACACC | |
| **gRNA4** | GGTA1 | f | caccGGTCTTCCAAGACGAGTTCG | |
| **gRNA4** | GGTA1 | r | aaacCGAACTCGTCTTGGAAGACC | |
| **BJO261** | CMAH PCR | f | CCCACCTCACTGAACATGCT | 413 |
| **BJO262** | CMAH PCR | r | AATCGGCCTCCAATTGAGCA | |
| **GL1213** | GGTA1 PCR | r | GGCCTATGTGATAATCCCAGCA | 520 |
| **GL1214** | GGTA1 PCR | f | AGGATGCCTTTGATAGAGCTGG | |
| **BJO280** | CMAH off target 1 | f | GAGGAAAAACAGCACTGGCTC | 278 |
| **BJO281** | CMAH off target 1 | r | GGACCCACTGGACATTCACA | |
| **BJO282** | CMAH off target 2 | f | CCGAAAGCTCCATAGAGACCC | 375 |
| **BJO283** | CMAH off target 2 | r | ATGCTACCTGGCACAAGTCT | |
| **BJO284** | CMAH off target 3 | f | CGCAGTGGTCTCCATTCAGT | 152 |
| **BJO285** | CMAH off target 3 | r | CTACTAACGCTGACAGTGGCT | |
| **BJO274** | GGTA1 off target 1 | f | CCAAGAACTCGGAGGCAGAC | 170 |
| **BJO275** | GGTA1 off target 1 | r | AGGCTGGAGGGTCAAAAAGT | |
| **BJO276** | GGTA1 off target 2 | f | AAGAAAGATCCTTGCACCACCT | 310 |
| **BJO277** | GGTA1 off target 2 | r | GTTCCGCGCATCCCATAGA | |
| **BJO286** | GGTA1 off target 3 | f | CTTCAGCTTGCACACTGGGT | 356 |
| **BJO287** | GGTA1 off target 3 | r | CTGGGGTGGAGCCTAGTTTTT | |
| **GL465** | Cas9 plasmid CBh | r | GGGCAGTTTACCGTAAATAC | 328 |
| **GL1358** | Cas9 insertion | f | CAGAGCTTCATCGAGCGGAT | 466 |
| **GL1359** | Cas9 insertion | r | CGAACAGGTGGGCATAGGTT | |
| **GL1041** | Plasmid sequencing | | ACTATCATATGCTTACCGTAAC | |
| **M13_f** | Plasmid sequencing (Massey) | f | CCCAGTCACGACGTTGTAAAACG | |

Plasmid overhangs indicated with lowercase letters. Underlined primers used for sequencing.

**Table S2. (a) Isolation and (b) mutation profiles of OFF3 and OFF4 double edited strains.**

a

| Parental cell line | Plated | First passage (%)[1] | Frozen (%)[2] | Mutation profiles |
|---|---|---|---|---|
| OFF3 | 107 | 77 (72%) | 61 (57%) | 12 |
| OFF4 | 90 | 83 (92%) | 16 (55%) | 14 |

[1] Normalised on cells plated.

[2] Cryopreserved strains normalised on uncontaminated cells plated.

b

| Cell line | Mutation profile | CMAH Edit 1 | CMAH Edit 2 | CMAH Edit 3 | GGTA1 Edit 1 | GGTA1 Edit 2 | GGTA1 Edit 3 | Individual strains isolated | Clonal strain | Frameshift both genes | Cas9 present | Off target sequence | SCT |
|---|---|---|---|---|---|---|---|---|---|---|---|---|---|
| OFF3 | A | +1 bp | | | +646 bp[A] | | | 1 | Yes | Yes | | | |
| | B | -27 bp | | | +1 bp | | | 3 | Yes | | No | | |
| | C | -3 bp | | | +1 bp | | | 2 | Yes | | No | | |
| | D | -1 bp | | | -1 bp | -5 bp | | 11 | Yes | Yes | No | WT | #11 |
| | E | -18 bp | | | -6 bp | -20 bp | | 1 | Yes | | ND | | |
| | F | -2 bp | | | +1 bp | -1 bp | | 1 | Yes | Yes | Yes | | |
| | G | -1 bp | -2 bp | | +1 bp | | | 3 | Yes | Yes | ND | WT | |
| | H | -1 bp | -2 bp | | +1 bp -1 bp[B] | | | 1 | Yes | | ND | | |
| | I | -1 bp | -4 bp | | -20 bp | | | 27 | Yes | Yes | ND | WT | |
| | J | +1 bp | -5 bp | | +1 bp | | | 3 | Yes | Yes | No | WT | #66 |
| | K | -21 bp | -44 bp | | +1 bp | -141 bp | | 1 | Yes | | No | | |
| | M | +1 bp | -1 bp | -5 bp | +1 bp | -1 bp | -5 bp | 1 | No[E] | Yes | | | |
| OFF4 | A | +1 bp | -15 bp | -22 bp | -8 bp | WT | | 1 | No | | Yes | | |
| | B | -3 bp | | | WT | +1 bp | | 1 | Yes | | Yes | | |
| | C | +1 bp | -3 bp | | +1 bp | -2 bp | | 1 | Yes | | No | | |
| | D | -1 bp | -8 bp | | -1 bp | -2 bp | | 1 | Yes | Yes | No | WT | #56 |
| | E | -3 bp | | | +1 bp | -2 bp | | 2 | Yes | | Yes | | |
| | F | +1 bp | +2 bp | | -1 bp | -15 bp | | 1 | Yes | | No | | |
| | G | -7 bp | ±81 bp[C] | | -85 bp | | | 1 | Yes | Yes | Yes | | |
| | H | -1 bp | -4 bp | | -1 bp | -14 bp | WT | 1 | No | | No | | |
| | I | -1 bp | -4 bp | | -1 bp | -3 bp | | 2 | Yes | | No | | |
| | J | -1 bp | -2 bp | | +1 bp | -1 bp | -2 bp | 1 | No | Yes | Yes | | |
| | K | -83 bp | -83 bp[D] | | +1 bp | +365 bp[A] | | 1 | Yes | Yes | No | | |
| | L | -4 bp | -6 bp | | +1 bp | -2 bp | | 1 | Yes | | No | | |
| | M | -5 bp | +2 bp | +3 bp | +1 bp | | | 1 | No | | No | | |
| | N | -5 bp | | | +1 bp | -12 bp | | 1 | Yes | | No | | |

[A] Inserted sequence matches part of Cas9 plasmid.
[B] Edits on same allele with 30 bp between them, suppressing frameshift mutation.
[C] 81 bp modified. Deletion of 37 bp along with an insertion of 44 bp roughly matching regions within CMAH.
[D] 3 SNPs not present on other allele.
[E] Likely mix of profiles D and J.
Cas9 plasmid insertion and off-target screening reported for strains tested. Shaded rows indicate strains chosen for SCT. ND, not definitive; WT, sequences in strain matched WT for that cell line.

**Table S3. Outcome of all *CMAH*$^{-/-}$ *GGTA1*$^{-/-}$ cloned lambs that survived past 115 days of gestation.**

| Year | Ewe ID | Lamb ID | Strain | Ewe steroid treatment | Delivery | Gestation length | Lamb outcome | Birth weight (kg) |
|---|---|---|---|---|---|---|---|---|
| 2018 | 605 | 2018.L02 | OFF3 #11 | 2x 20mg Dex | Non-assisted natural | 151 | Died <24hr | 3.6 |
| 2018 | 3340 | 2018.L03 | OFF3 #66 | 2x 20mg Dex | Surgical recovery | 149 | Euthanised <1hr | 6.3 |
| 2018 | 627 | 2018.L04 | OFF3 #66 | 2x 20mg Dex | Surgical recovery | 150 | Euthanised <1hr | 2.2 |
| 2018 | 606 | 2018.L05 | OFF3 #66 | None | Non-assisted natural | 144 | Died <24hr | 3.7 |
| 2019 | 608 | 2019.L02 | OFF3 #66 | 5 mg TMA/ 20mg Dex | Surgical recovery | 147 | Died <1hr | 3.0 |
| 2019 | 601 | 2019.L03 | OFF4 #56 | 5 mg TMA/ 20mg Dex | Surgical recovery | 147 | Died <1hr | 3.5 |
| 2019 | 637 | 1917 | OFF4 #56 | 5 mg TMA/ 20mg Dex | Surgical recovery | 148 | Alive | 6.4 |
| 2019 | 3362 | 1918 | OFF4 #56 | 5 mg TMA/ 2x 20mg Dex | Surgical recovery | 148 | Alive | 3.0 |
| 2019 | 691 | 2019.L05 | OFF3 #66 | 5 mg TMA/ 2x 20mg Dex | Surgical recovery | 147 | No HB at birth | 3.8 |
| 2019 | 635 | 2019.L06 | OFF4 #56 | 5 mg TMA/ 6mg Dex/2x 20mg Dex | Surgical recovery | 148 | No HB at birth | 5.7 |
| 2019 | 619 | 2019.L07 | OFF4 #56 | 5 mg TMA/ 6mg Dex/2x 20mg Dex | Surgical recovery | 148 | Died <1hr | 5.4 |
| 2019 | 619 | 1919 | OFF4 #56 | 5 mg TMA/ 6mg Dex/2x 20mg Dex | Surgical recovery | 148 | Alive | 3.3 |
| 2019 | 489 | 1920 | OFF4 #56 | 5 mg TMA/ 6mg Dex/2x 20mg Dex | Surgical recovery | 146 | Alive | 3.9 |
| 2019 | 493 | 1921 | OFF4 #56 | 5 mg TMA/ 6mg Dex/2x 20mg Dex | Surgical recovery | 146 | Alive | 3.5 |
| | | | | **Aborted** | | | | |
| 2018 | 3352 | 2018.L01 | OFF3 #11 | None | Non-assisted natural | 138 | Aborted | NA |
| 2019 | 9149 | 2019.L01 | OFF4 #66 | None | Non-assisted natural | 119 | Aborted | NA |
| 2019 | 601 | 2019.L04 | OFF4 #56 | 5 mg TMA/20mg Dex | Surgical recovery | 147 | No HB at birth | 1.8 |
| 2019 | 648 | - | OFF4 #56 | 5 mg TMA/6mg Dex/2x 20mg Dex | Surgical recovery | 145 | Aborted | 2.0 |

**Table S4. Post mortem findings from *CMAH*$^{-/-}$ *GGTA1*$^{-/-}$ cloned lambs that died within first 24 hrs.**

| Lamb ID | Strain | Birth weight (kg) | Lamb outcome | Musculoskeletal | Pulmonary | Hepatic | Renal | Intestinal | Cardiovascular |
|---|---|---|---|---|---|---|---|---|---|
| 2018.L02 | OFF3 #11 | 3.6 | Found dead (Dystocia) | Moderate brachygnathism | Pulmonary atelectasis | NSF | Mild hydronephrosis | NSF | NSF |
| 2018.L03 | OFF3 #66 | 6.3 | Euthanised < 1hr | Moderate brachygnathism | Pulmonary atelectasis | NSF | Mild hydronephrosis | Mesenteric twist | NSF |
| 2018.L04 | OFF3 #66 | 2.2 | Euthanised < 1hr | Mild brachygnathism; Rotated contracted limbs | Pulmonary atelectasis; Pulmonary hamartoma | NSF | Mild hydronephrosis | NSF | NSF |
| 2018.L05 | OFF3 #66 | 3.7 | Found dead (Dystocia) | Mild brachygnathism; Signs of severe dystocia (Head/neck/eye/tongue swollen. Bloody mucoid discharge around nares) | Pulmonary atelectasis | Mild cholestasis | Moderate hydronephrosis | NSF | Patent foramen ovale |
| 2019.L02 | OFF3 #66 | 3.0 | Died < 1hr | NSF | Pulmonary atelectasis | Moderate cholestasis | Mild hydronephrosis | Hypoplasia of the distal small intestine, caecum and colon; Dilation of the proximal small intestine. | Extramedullary haematopoiesis |
| 2019.L03 | OFF4 #56 | 3.5 | Died < 1hr | Mild brachygnathism | Pulmonary atelectasis; pulmonary hamartoma | Marked hepatic iron storage; cholestasis | Severe hydronephrosis; renal cysts | NSF | NSF |
| 2019.L05 | OFF3 #66 | 3.8 | No HB at birth | Mild brachygnathism | Pulmonary atelectasis | NSF | Moderate hydronephrosis | Dysplasia of the spiral colon | NSF |
| 2019.L06 | OFF4 #56 | 5.7 | No HB at birth | Generalized muscular hypertrophy; contracted forelimbs | Pulmonary atelectasis | NSF | Moderate hydronephrosis | NSF | NSF |
| 2019.L07 | OFF4 #56 | 5.4 | Died < 1hr | NSF | Pulmonary atelectasis | NSF | Moderate hydronephrosis | NSF | NSF |

NSF, non-significant findings.

**Table S5. Breeding of *CMAH*$^{-/-}$ *GGTA1*$^{-/-}$ cloned ewes.**

| Year | Method | Genotype (*CMAH* & *GGTA1*) | | | | Pregnancy |
|---|---|---|---|---|---|---|
| | | Dam | Sire | Embryo | Surrogate | |
| **2021** | Natural mating | DKO | WT | -/+ & -/+ | DKO | No (2 cycles)[A] |
| **2022** | OPU/IVF/ET | DKO | WT | -/+ & -/+ | WT | Yes (2 lambs) |
| **2023** | Natural mating | DKO | WT | -/+ & -/+ | DKO | No (2 cycles)[A] |

[A] Females were naturally bred to a ram for two consecutive estrus cycles. All contemporary WT controls (n=3) became pregnant and lambed without incident.
DKO, double knockout for CMAH & GGTA1; OPU/IVF/ET, ovum pick-up, in vitro fertilisation, and embryo transfer; WT, wild-type.

**Table S6. OPU results for *CMAH*$^{-/-}$ *GGTA1*$^{-/-}$ and wild-type (WT) cloned ewes.**

| Genotype | n[1] | N[2] | Follicles aspirated | Oocytes recovered | Follicles/donor | Oocytes/donor | Oocytes/follicle |
|---|---|---|---|---|---|---|---|
| DKO | 2 | 5 | 126 | 120 | 12.6 | 12.0 | 1.0 |
| WT | 2–4[A] | 4 | 166 | 153 | 11.9 | 10.9 | 0.9 |

[1] Number of OPU runs.

[2] Number of ewes per run.

[A] 1 ewe was used twice, remaining 3 ewes were used 4 times.

DKO, double knockout for CMAH & GGTA1; WT, wild-type.

**Table S7. Development of IVF embryos derived from *CMAH*$^{-/-}$ *GGTA1*$^{-/-}$ or wild-type (WT) oocytes.**

| Oocyte genotype | n[1] | nIVC[2] | >1cell (%)[3] | B 1–3 (%)[4] | B 1–2 (%)[5] |
|---|---|---|---|---|---|
| DKO | 1 | 48 | 46 (96) | 10 (22) | 6 (60) |
| WT | 2 | 57 | 51 (89) | 16 (31) | 12 (75) |

[1] Number of OPU/IVF runs.
[2] Number of embryos placed into in vitro culture (IVC).
[3] Cleavage normalised on nIVC.
[4] Total blastocyst development normalised on > 1-cell embryos.
[5] Proportion of grade 1–2 blastocysts normalised on total number of blastocysts.
DKO, Double knockout for CMAH & GGTA1; WT, wild-type.

**Table S8.** ***In vivo*** **survival of embryos derived from** ***CMAH***$^{-/-}$ ***GGTA1***$^{-/-}$ **or wild-type (WT) oocytes.**

| Oocyte genotype | Recipient WT ewes | Embryos transferred | Lambs born (%)[1] | Lambs alive (%)[1] |
|---|---|---|---|---|
| DKO | 3 | 9 | 2 (22) | 0 (0) |
| WT | 12 | 36 | 11 (31) | 7 (19) |

[1] Normalised on blastocysts transferred into WT recipient ewes.
DKO, Double knockout for CMAH & GGTA1; WT, wild-type.
